# Inhibitory and *in silico* molecular docking of *Xeroderris stuhlmannii* (Taub.) Mendonca & E.P. Sousa phytochemical compounds on human α-glucosidases

**DOI:** 10.1101/2022.09.16.508336

**Authors:** Brilliant Nyathi, Jonathan Tatenda Bvunzawabaya, Chido Venissa P Mudawarima, Emily Manzombe, Kudakwashe Tsotsoro, Major Allen Selemani, Gadzikano Munyuki, Freeborn Rwere

## Abstract

**Ethnopharmacological relevance:** Herbal traditional medicine is used by millions of people in Africa for treatment of ailments such as diabetes mellitus, stomach disorders and respiratory diseases. *Xeroderris stuhlmannii (Taub.*) Mendonca & E.P. Sousa (*X. stuhlmannii* (Taub.)) is a medicinal plant used traditionally in Zimbabwe to treat type 2 diabetes mellitus (T2DM) and its complications. However, there is no scientific evidence to support its inhibitory effect against digestive enzymes (α-glucosidases) that are linked to high blood sugar in humans.

**Aim of the study:** : This work aims to investigate whether bioactive phytochemicals of crude *X. stuhlmannii* (Taub.) can scavenge free radicals and inhibit α-glucosidases in order to reduce blood sugar in humans.

**Materials and methods:** Here we examined the free radical scavenging potential of crude aqueous, ethyl acetate and methanolic extracts of *X. stuhlmannii* (Taub.) using the diphenyl-2-picrylhydrazyl assay *in vitro*. Furthermore, we carried out *in vitro* inhibition of α-glucosidases (α-amylase and α-glucosidase) by the crude extracts using chromogenic 3,5-dinitrosalicylic acid and p-nitrophenyl-α-D-glucopyranoside substrates. We also used molecular docking approaches (Autodock Vina) to screen for bioactive phytochemical compounds targeting the digestive enzymes.

**Results:** Our results showed that phytochemicals in *X. stuhlmannii* (Taub.) aqueous, ethyl acetate and methanolic extracts scavenged free radicals with IC_50_ values ranging from 0.002-0.013 μg/mL. Furthermore, crude aqueous, ethyl acetate and methanolic extracts significantly inhibited α-amylase and α-glucosidase with IC_50_ values of 10.5-29.5 μg/mL (versus 54.1±0.7 μg/mL for acarbose) and 8.8-49.5 μg/mL (versus 161.4±1.8 μg/mL for acarbose), respectively. *In silico* molecular docking findings and pharmacokinetic predictions showed that myricetin is likely a novel plant-derived α-glucosidase inhibitor.

**Conclusion:** Collectively, our findings suggest pharmacological targeting of digestive enzymes by *X. stuhlmannii* (Taub.) crude extracts may reduce blood sugar in humans with T2DM via inhibition of α-glucosidases.

## INTRODUCTION

Diabetes mellitus is a major epidemic of the 21^st^ century that affects millions of people worldwide and contributes to high morbidity and mortality in adult populations (Lin et al., 2020; Saeedi et al., 2019; Sun et al., 2022). There are two major forms of diabetes mellitus: type 1 and type 2 diabetes mellitus (T2DM). T2DM is the most common form, and affects 462 million people (about 6.28% of the world’s population) (Khan et al., 2019). In Africa, there is an increase in the incidence of T2DM in urban areas because of rural to urban migration and lifestyle changes (Godman et al., 2020). In the case of Zimbabwe, it was estimated that approximately 5.7% of its population is living with T2DM (Mutowo et al., 2015). Since the cost of healthcare is very high in Zimbabwe, many rural and urban people are using herbal medicines for the treatment and management of diabetes mellitus and its complications (Mutowo et al., 2016).

One effective therapeutic approach of reducing postprandial glucose in T2DM patients involves inhibiting α-glucosidase enzymes with drugs such as acarbose, miglitol and voglibose (Vieira et al., 2019). These α-glucosidase inhibitors reduce intestinal glucose absorption by delaying carbohydrate digestion (Vieira et al., 2019). However, a number of side effects have been reported in patients using α-glucosidase inhibitors such as acarbose and voglibose (Zhang et al., 2016). This has led to the search for new plant derived natural product inhibitors targeting digestive enzymes with fewer side effects to reduce postprandial glucose in T2DM patients.

For many decades, herbal traditional medicines have been clinically used to treat metabolic diseases such as diabetes mellitus (Nyakudya et al., 2020; Usai et al., 2022). Phytochemical compounds in plants such as polyphenols and flavonoids have antioxidative properties, and are capable of scavenging free radicals and reduce oxidative stress, which in-turn lowers blood glucose (Sarker and Oba, 2020; Tan et al., 2018). In addition, the phytochemical compounds in plant extracts improve insulin resistance and glucagon production by binding to β-cells and α-cells in response to high blood glucose (Ansari et al., 2022; Williamson and Sheedy, 2020). Additionally, phytochemical compounds reduce glucose absorption in the gastrointestinal tract by inhibiting digestive enzymes such as α-amylase and α-glucosidase (Shakoor et al., 2020; Sun et al., 2020). High blood glucose is associated with many comorbidities such as heart disease, stroke, kidney disease, eye problems (diabetic retinopathy, glaucoma or cataracts), dental disease, peripheral neuropathy and feet problems (peripheral artery disease, PAD) that reduce the quality of life and life expectancy of diabetic patients (Forbes and Cooper, 2013). However, despite the widespread use of medicinal plants by many African people to treat diabetes mellitus, many plants native to Zimbabwe such as *X. stuhlmannii* (Taub.) have not received much attention to assess their *in vitro* inhibitory effect on digestive enzymes.

*X. stuhlmannii* (Taub.) from *Xeroderris* Roberty genus of the Fabaceae family is a medicinal plant widely used traditionally to treat many ailments across Zimbabwe (Selemani et al., 2021). Different parts of the plant are used to treat diabetes mellitus, bacterial wound infections, coughs, diarrhea, malaria, colds, rheumatoid arthritis, stomachache, dysentery and eye infections (Asase et al., 2005; Bello et al., 2019; Chinemana et al., 1985; Shopo et al., 2022). *In vitro* antibacterial studies showed that the bark extracts effectively fight bacterial pathogens that cause gastrointestinal disorders in humans (Selemani et al., 2021). Ethnobotanical survey of *X. stuhlmannii* (Taub.) in the Central Region of Togo showed that leaves and root decoctions are used to treat diabetes mellitus (Karou et al., 2011) as well as female sexual disorders and infertility (Tchacondo et al., 2011). However, they were also associated with side effects such as diarrhea and polyuria (Tchacondo et al., 2011).

Spectrochemical characterization of *X. stuhlmannii* (Taub.) by gas chromatography mass spectroscopy (GC MS) and liquid chromatography tandem mass spectroscopy (LC MS/MS) confirmed the presence of thirty-six (36) important bioactive phytochemical compounds categorized into phenolic, flavonoid and alkaloid compounds (Selemani et al., 2021). Whether some of the phytochemical compounds present in *X. stuhlmannii* (Taub.) bark extracts have inhibitory effects on digestive enzymes remains unknown. In Zimbabwe, decoctions of *X. stuhlmannii* (Taub.) barks and roots are traditionally used to treat T2DM and its complications. However, no scientific evidence supports its use to treat T2DM. For this reason, this study aims to determine the inhibitory effects of bark and root extracts of *X. stuhlmannii* (Taub.) on α-amylase and α-glucosidase because their activity may lead to increased blood sugar in humans.

## MATERIALS AND METHODS

### Collection of plant materials and preparation of *X. stuhlmannii* (Taub.) root and bark extracts

*X. stuhlmannii* (Taub.) root and bark samples were obtained from a wild tree and the voucher specimen of the plant is *Luckmore Kazingizi Number 1*. The specimen is stored at National Herbarium & Botanic Garden in Harare, Zimbabwe. Plant samples were pre-washed with running tap water and rinsed with distilled water, followed by sun drying for several days at approximately 25°C. The dried samples were grinded and sieved with a 1 mm sieve to obtain finely divided powdered samples. Crude root and bark extracts were obtained by the maceration method from dried plant extracts (5 g) using 5 mL of solvents (water, methanol (Glassworld, South Africa), methanol/water (1:1), ethyl acetate (Glassworld, South Africa) or ethyl acetate/water (1:1)) and filtered through (Whatman No. 1) filter paper. Following extraction, the solvents were removed using a rotary evaporator and further dried in a fumehood for a few days. The percentage yield of plant extracts was recorded after completely drying the samples (see Table 1).

**Table 1.** Extraction yields, total phenolic and flavonoid contents of *X. stuhlmannii* (Taub.) roots and bark extracts.

| Plant species | Plant part used | Solvent used | Yield of the plant (%w/w) | Total<sup>a</sup> phenolic content (mg GAE/g DW) | Total<sup>a</sup> flavonoid content (mg QE/g DW) |
| --- | --- | --- | --- | --- | --- |
| *Xeroderris stuhlmannii* (Taub) | Root | Methanol (100%) | 09.4 | 11.5±0.1 | 15.6±0.2 |
|  |  | Methanol/water (50:50%) | 07.5 | 27.7±0.2 | 33.6±0.5 |
|  |  | Ethyl acetate (100%) | 04.2 | 25.7±0.1 | 21.3±0.4 |
|  |  | Ethyl acetate/water (50:50%) | 06.5 | 58.1±0.6 | 58.1±1.2 |
|  |  | Water (100%) | 11.0 | 35.0±0.3 | 50.9±0.8 |
|  | Bark | Methanol (100%) | 13.6 | 07.5±0.1 | 17.4±0.2 |
|  |  | Methanol/water (50:50%) | 08.3 | 16.7±0.2 | 20.2±0.3 |
|  |  | Ethyl acetate (100%) | 05.2 | 19.6 ±0.1 | 19.4±0.5 |
|  |  | Ethyl acetate/water (50:50%) | 04.5 | 80.1±0.8 | 58.8±1.8 |
|  |  | Water (100%) | 11.8 | 35.5±0.3 | 44.9±0.4 |

<sup>a</sup>n=3
GAE/g DW is gallic equivalence per gram of dry weight
QE/g DW is quercetin equivalents per gram of dry weight

### Determination of total phenolic content

The total phenolic concentration was determined spectrophotometrically using the Shimadzu UV-1900i spectrophotometer, (Shimadzu, Japan) according to the Folin-Ciocalteu method (Mbhele et al., 2022). Briefly, 0.2 mL of crude plant extract was added to 0.2 mL of 10% methanol, followed by 5 mL of 10% Folin-Ciocalteau phenol reagent (HiMedia Laboratories, India) and 4.0 mL of 7.5% Na_2_CO_3_ (Associated chemical enterprises, South Africa). The mixture was incubated at 25°C in the dark for 30 min. After 30 min, absorbance of the mixture was determined at 765 nm. Total phenolics were quantified by a calibration curve obtained from measuring the absorbance of known concentrations of gallic acid (Sigma-Aldrich, USA) standard. All measurements were done in triplicates and on two separate days to check the reproducibility of results. The data is expressed as mg of gallic acid equivalents/g of dry extract (mg GA/g DE).

### Determination of total flavonoid content

The total flavonoid content in *X. stuhlmannii* (Taub.) was determined spectrophotometrically using the aluminum chloride (AlCl_3_) (Associated chemical enterprises, South Africa) colorimetric assay according to published procedures (Mbhele et al., 2022). Briefly, 0.15 mL of plant extract was mixed with 0.45 mL methanol and 0.6 mL of 2% aluminum chloride. After mixing, the solution was incubated for 60 min at 25°C in the dark, followed by absorbance measurement at 420 nm. Quercetin dihydrate (Fisher Scientific, USA) was used as a standard for the calibration curve. The standard solutions of quercetin were prepared by serial dilutions using methanol. Total flavonoids content of the extract is expressed as mg quercetin equivalents (QE) per dry extract (mg/g).

### *In vitro* antioxidant activity of *X. stuhlmannii* (Taub.) extracts

Free radical scavenging activity of *X. stuhlmannii* (Taub.) crude extracts and garlic acid were determined *in vitro* by diphenyl-2-picrylhydrazyl (DPPH) assay according to published procedures with a slight modification (Mbhele et al., 2022). Briefly, 1 mL of 0.1 mM DPPH (Research Lab Fine Chem Industries, India) in methanol was added to 3 mL of various concentrations (0.0005-0.03 μg/mL) of plant extracts at 25°C. The samples were vigorously mixed and incubated in the dark for 30 min. After 30 min, absorbance of plant samples was measured at 517 nm. Measurements were carried out in triplicates and on two separate days for reproducibility.

The percentage (%) DPPH free scavenging activity (RSA) was calculated as presented in equation 1, below and the IC_50_ values denote the concentration of the sample required to scavenge 50% DPPH free radicals. Gallic acid (Sigma-Aldrich, South Africa) was used as a positive control.

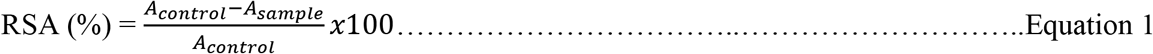

Where, *A_sample_* is the absorbance of DPPH and crude extract; *A_control_* is the absorbance of DPPH without crude extract.

### Inhibition of α-amylase by crude *X. stuhlmannii* (Taub.) extracts

The inhibition of α-amylase by crude root and bark extracts of *X. stuhlmannii* (Taub.) was performed using the 3,5-dinitrosalicylic acid (DNSA) as reported elsewhere (Kifle et al., 2021). Briefly, crude root and bark extracts of *X. stuhlmannii (Taub.*) were dissolved in 20 mM sodium phosphate buffer, pH 6.9 to give concentrations ranging from 0 to 100 μg/mL. Bacterial α-amylase (0.25 ml, 2 units/mL) (Phillip Harris Manufacturing Ltd, United Kingdom) was mixed with 0.25 mL sample extract (0-100 μg/mL) and incubated for 10 min at 30 °C. After 10 min, 0.25 mL (1% w/v) of the starch (Associate chemical enterprises, South Africa) solution was added to each tube and incubated for a further 3 min. The reaction was terminated by adding 0.25 mL DNSA reagent (12 g of sodium potassium tartrate tetrahydrate (Associated chemical enterprises, South Africa) in 8.0 mL of 2 M NaOH (Glassworld, South Africa) and 20 mL of 96 mM 3,5-DNSA (Sisco Research Laboratories, India), and boiled for 10 min in a water bath at 85°C. The mixture was cooled to 25°C and diluted with 5 mL of distilled water. Absorbance was immediately measured at 540 nm using a UV-visible spectrophotometer. The control experiment was carried out with enzyme and starch solution only. Acarbose (Fisher Scientific, USA) was used as a positive control. The percentage (%) inhibition was calculated as follows:

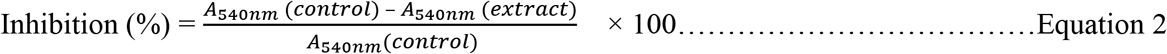

### Inhibition of α-glucosidase by crude *X. stuhlmannii* (Taub.) extracts

The α-glucosidase inhibitory activity was measured using p-nitrophenyl-α-d-glucopyranoside (pNPG) (Sigma-Aldrich, USA) and various concentrations of *X. stuhlmannii* (Taub.) crude extracts according to published procedures (Kifle et al., 2021). Briefly, 50 μL of crude root and bark extracts at varying concentrations (5-400 μg/mL) was mixed with 10μL of α-glucosidase from Aspergillus, Niger (Sigma-Aldrich, South Africa) (1 U/mL) and 120 μL of 20 mM sodium phosphate buffer, pH 6.9. The resultant mixture was incubated at 37 °C for 20 min. After 20 min, the reaction was initiated by adding 20 μL of 1M pNPG and the resultant mixture was incubated for 30 min at 25°C. The reaction was terminated by addition of 0.1M of Na_2_CO_3_ (50 μL) and further diluted with 1.5 mL of 20 mM sodium phosphate buffer, pH 6.9. Absorbance of the mixture was immediately measured at 405 nm. Acarbose (Fisher Scientific, USA) was used as a positive control. The percentage (%) inhibition was calculated as follows:

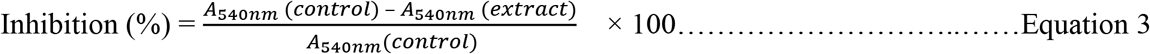

### Molecular docking

Molecular docking of phytochemical compounds was carried out on human lysosomal acid-α-glucosidase, hGAA (PDB: 5NN8) and human pancreatic α-amylase, HPA (PDB: 2QV4) (Maurus et al., 2008; Roig-Zamboni et al., 2017). The X-ray crystallographic structures of hGAA and HPA were retrieved from the protein data bank (https://www.rcsb.org/), visualized and prepared for docking using Bovia Discovery studio visualizer v21.1.0.20298. Prior to docking, co-crystallized water molecules and non-essential small organic molecules were removed from the crystal structures. The retained protein-ligand complexes were then protonated, optimized and typed using Charmm and MMFF94 forcefields.

A dataset of thirty-six (36) phytochemical compounds identified using gas chromatography mass spectroscopy (GC MS) and liquid chromatography with tandem mass spectroscopy (LC MS/MS) from *X. stuhlmannii* (Taub.) was used for molecular docking (Selemani et al., 2021). The structures of *X. stuhlmannii* (Taub.) were retrieved from PubChem database (https://pubchem.ncbi.nlm.nih.gov/) in SDF format, protonated using Discovery Studio visualizer and energetically optimized with MMFF94 forcefield using RDkit optimizer node in KNIME Analytics v4.3.3 (Berthold et al., 2008). Molecular docking simulations were performed using Autodock Vina (Vina) in Pyrx v0.8 (Dallakyan and Olson, 2015; Trott and Olson, 2009). Co-crystalized acarbose ligand (positive control) in human lysosomal acid-α-glucosidase and pancreatic α-amylase was used for validating the docking procedures. The docking scores from Autodock Vina are reported as binding affinity (kcal/mol). Discovery studio visualizer was used to analyze the binding interactions of the docked protein-ligand complexes.

### The physicochemical, drug-likeness, medicinal, ADME and toxicity properties of *X. stuhlmannii* (Taub.) compounds

The physicochemical properties (molecular weight (MW), hydrogen bond donor count (HBD), hydrogen bond acceptor count (HBA), rotatable bond count (RB), lipophilicity and water solubility) were computed using SwissADME (http://www.swissadme.ch/, accessed on 04/07/2022). Additionally, the drug-likeness, oral bioavailability and medicinal chemistry properties of the phytochemical compounds were evaluated using SwissADME. Finally, the absorption, distribution, metabolism, and elimination (ADME) were computed using ADMElab 2.0 (accessed on 05/08/2022).

### Statistical analysis

All experiments were performed in triplicates and on two separate days. Data analysis was performed by one-way analysis of variance (ANOVA) using GraphPad Prism version 8.0. The results are expressed as means of three replicate determinations ± standard deviation. The IC_50_ values were obtained with GraphPad Prism version 9.5.1 using non-linear regression fit in which the percentage of inhibition (% inhibition) was plotted against the concentration of extracts with four variables.

## RESULTS

### Crude extract yields, total phenolic and total flavonoid content of *X. stuhlmannii* (Taub.) root and bark extracts

The percentage (%) yield of crude root and bark *X. stuhlmannii* (Taub.) extracts are summarized in Table 1. The yield of plant extracts ranged from 4.2% to 13.6% for bark and roots. Methanol and water extracted more secondary plant metabolites/phytochemicals compared to ethyl acetate. The quantified total phenolics (TFC) of bark and root extracts significantly varied (7.5-80.1 gallic acid equivalent per g of dry weight) between methanol, ethyl acetate and water (Table 1). Aqueous ethyl acetate and water extracted more polyphenols compared to all other solvents, whereas methanol extracts had the lowest polyphenolic content.

The total flavonoid content (TFC) of bark and root extracts of *X. stuhlmannii* (Taub.) are expressed as mg quercetin equivalents/g of dry weight extract (Table 1). The total flavonoid content of the root extract and bark extracts ranged from 15.6-58.8 quercetin equivalent per g of dry weight. Aqueous bark ethyl acetate extract had the highest flavonoid content (58.8±0.8 mg of QE/g dry weight). The flavonoid content of bark and root extracts were significantly different from each other depending on the solvent used to extract phytochemicals.

### DPPH Free Radical Scavenging Activity of root and bark extracts

The *in vitro* antioxidant activity assay was carried out to assess the ability of crude *X. stuhlmannii* (Taub.) root and bark extracts to scavenge free radicals such as 2,2-di-(4-tert-octylphenyl)-1-picrylhydrazyl free radical (DPPH). Table 2 summarize the antioxidant results obtained for the crude methanolic and ethyl acetate root and bark extracts. Gallic acid was used as a positive control. As shown in Table 2, both root and bark extracts significantly scavenged DPPH radicals (IC_50_ values ranged from 0.002-0.013 μg/mL for methanolic and ethyl acetate extracts). The lower the IC_50_ value of a plant extract, the higher its antioxidant activity (Mbhele et al., 2022). Gallic acid displayed a good scavenging effect against the DPPH radical with a calculated IC_50_ value of 0.215 μg/mL. However, its inhibitory activity was moderately lower than that of root and bark extracts.

**Table 2.**
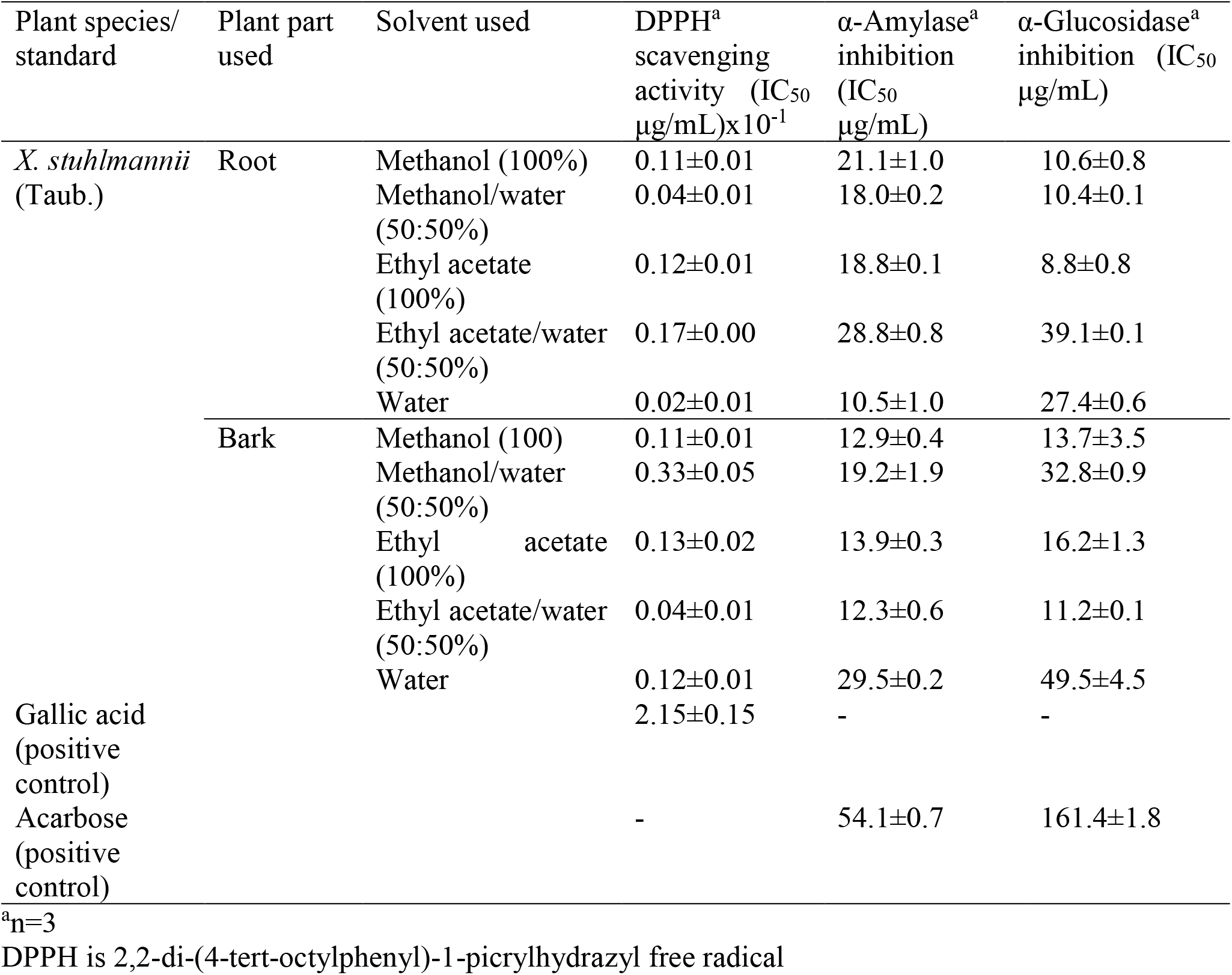
Concentration of *X. stuhlmannii* (Taub.) root and bark extracts that causes 50% inhibition (IC_50_) values in DPPH radical scavenging, α-amylase and α-glucosidase inhibitory assays.

### *In vitro* inhibitory effect of *X. stuhlmannii* (Taub.) crude extracts on α-amylase and α-glucosidase

Polyphenols in plants have antioxidant properties and are reported to exert anti-hyperglycemic effects through non-specific binding to glucose transporters and competitively inhibit digestive enzymes (Kim et al., 2016; Mala et al., 2022). The inhibition of α-amylase and α-glucosidase by *X. stuhlmannii* (Taub.) root and bark extracts are described in Table 2. *X. stuhlmannii* (Taub.) methanol, ethyl acetate and water root extracts inhibited α-amylase with IC_50_ values ranging from 10.5 to 28.8 μg/mL. Inhibition of α-amylase by methanol (100% solvent) and ethyl acetate (100% solvent) root extracts occurred with IC_50_ values of 21.1±1.0 μg/mL and 18.8±0.1 μg/mL, respectively. Aqueous methanol (IC_50_=18±0.2 μg/mL) and aqueous ethyl acetate (IC_50_=28.8±0.8 μg/mL) root extracts had modest inhibition of α-amylase. In contrast, acarbose inhibited α-amylase with an IC_50_ value of 54.4±6.4 μg/mL. The root water extracts significantly inhibited α-amylase compared to acarbose (IC_50_=10.5±1.0 μg/mL versus 54.4±6.4 μg/mL for acarbose). The IC_50_ values of bark extracts were comparable to those of root extracts, except for water (IC_50_=29.5 μg/mL for bark versus 10.5 μg/mL for root). These results show that root and bark extracts of *X. stuhlmannii* (Taub.) significantly inhibited α-amylase compared to the commercial antidiabetic drug, acarbose.

We next tested the *in vitro* inhibition of α-glucosidase by *X. stuhlmannii* (Taub.) methanol and ethyl acetate root and bark extracts. Inhibition of α-glucosidase by methanolic, ethyl acetate and water root extracts occurred with IC_50_ values ranging from 8.8 μg/mL to 49.5 μg/mL. The calculated IC_50_ values for methanolic (100%) and ethyl acetate (100%) root extracts were 10.6±0.8 μg/mL and 8.8±0.8 μg/mL. Only aqueous methanol (bark), aqueous ethyl acetate (root) and water (bark) extracts had higher IC_50_ values compared to other extracts but were significantly lower than that of acarbose. Water bark extract had the highest IC_50_ value of 49.5±4.5 μg/mL. In contrast, α-glucosidase inhibition by acarbose occurred with an IC_50_ value of 161.4±1.8 μg/mL.

### *Molecular docking of the phytochemical compounds of Xeroderris stuhlmannii* Taub

A recent publication by (Selemani et al., 2021) identified thirty-six (36) phytochemical compounds from bark extract of *X. stuhlmannii* (Taub.). To evaluate the inhibitory properties of the phytochemical compounds, we virtually screened them using Autodock Vina against human pancreatic α-amylase (HPA) and lysosomal acid-α-glucosidase (hGAA). The docking method was validated by redocking co-crystallized acarbose and an acarbose derivative to the active site of the enzymes. Figure 1 shows the graphical summary of binding affinities of selected phytochemical compounds and acarbose (positive control) against HPA and hGAA enzymes. The binding affinities of all thirty-six phytochemical compounds against HPA ranged from −10.3 to −5.6 kcal/mol, whereas those for hGAA ranged from −8.4 to −5.6 kcal/mol. Khasianine, brassinolide, oleanolic aldehyde and castasterone had higher binding affinities (more negative) than acarbose in HPA. Oleanolic aldehyde and apiin had higher binding affinities than acarbose in hGAA. Khasianine had the same binding affinity as acarbose in hGAA. Phytochemical compounds with higher or equal binding affinities than acarbose in both HPA and hGAA were considered for pharmacokinetic prediction studies. In addition, a few other compounds with lower binding affinities than acarbose such as ursolic acid, myricetin, and myricitrin were also included in pharmacokinetic studies because they are known to inhibit digestive enzymes (Castro et al., 2015; Kim et al., 2020; Williams et al., 2012).

**Figure 1.**
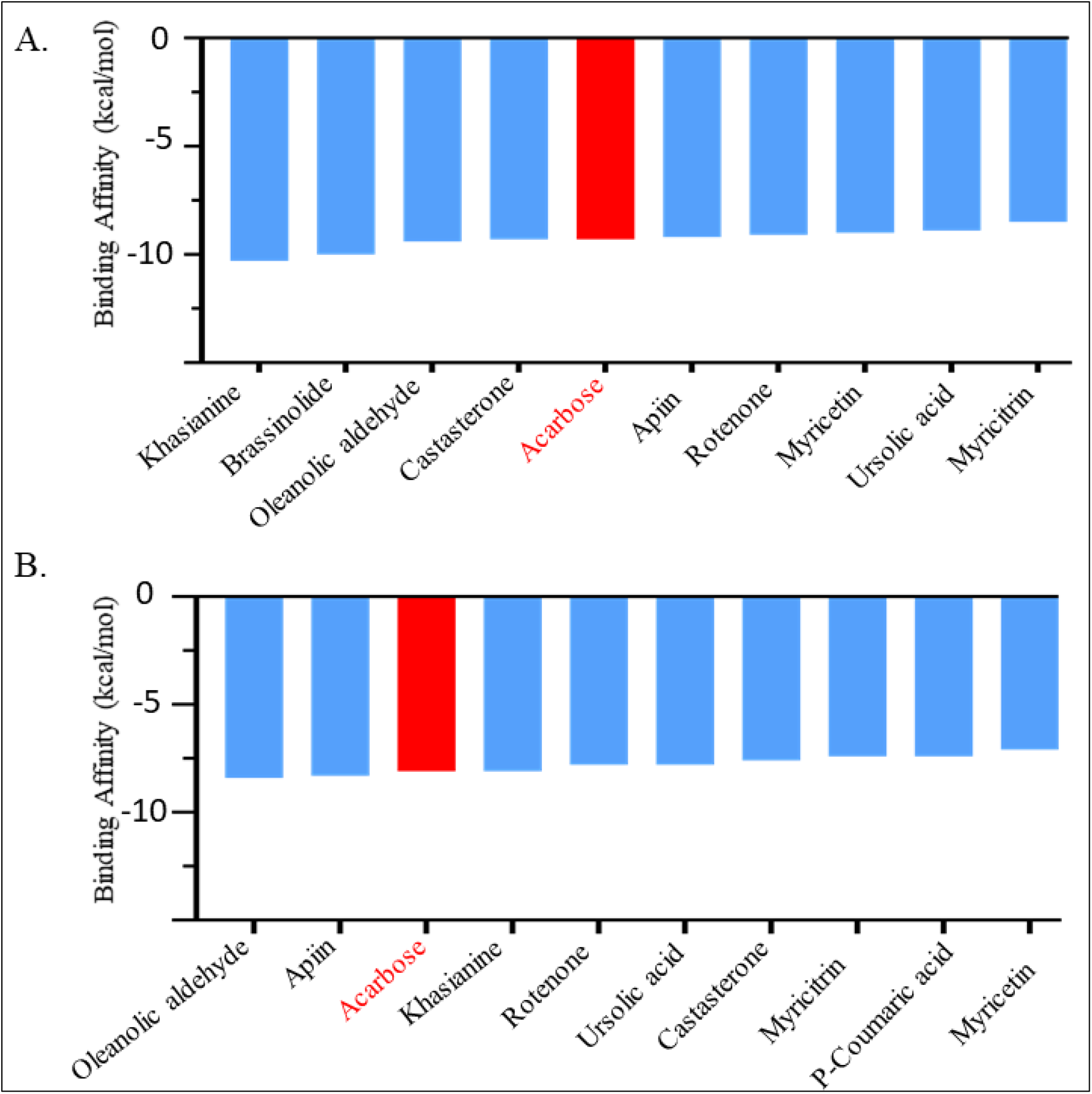
Variation in binding affinity of selected compounds of *X. stuhlmannii* (Taub.) and acarbose against: A. human pancreatic α-amylase (HPA) and B. human lysosomal acid-α-glucosidase (hGAA).

Intermolecular interactions between selected ligands (those with higher binding affinities than acarbose as well as those known to inhibit digestive enzymes) and the active site residues of HPA and hGAA are shown in Table 3 and Figures 2&3. The major intermolecular interactions observed between the ligands and digestive enzymes were hydrogen-bonding (H-bonding), π-π, electrostatic and hydrophobic interactions. The majority of selected compounds formed H-bonds with bond lengths less than 3 Å. When the co-crystallized acarbose derivative was redocked to the binding site of the human pancreatic amylase (HPA), the root mean square deviation (RMSD) was 1.62Å (Figure 2). In this orientation, the acarbose derivative formed two strong H-bonds with the catalytic residues, GLU233 and ASP300. In addition, the acarbose derivative was also stabilized by 10 extra H-bonds with neighboring amino acids (Figure 2). All other docked compounds, except myricetin formed a H-bond with at least one of the three key catalytic residues (ASP197, GLU233 and ASP300) of HPA (Figure 2). Brassinolide, khasianine and apiin formed more H-bonds than other phytochemical compounds leading to high binding affinities (Figure 2). To get insight into the interaction of hGAA with the phytochemical compounds, we docked all thirty-six phytochemical compounds into the active site of hGAA (Figure 1) and those with higher binding affinity than acarbose are shown in Figure 3. Acarbose is stabilized within the active site by four strong H-bonds with ASP282, ASP404, ASP600 and ASP616. When apiin was docked into the substrate binding site of hGAA, it was more stabilized by H-bonds compared to acarbose and other compounds with similar binding affinities (Figure 3). Oleanolic aldehyde did not form H-bonds with hGAA, whereas myricetin formed two H-bonds with hGAA (Figure 3).

**Figure 2.**
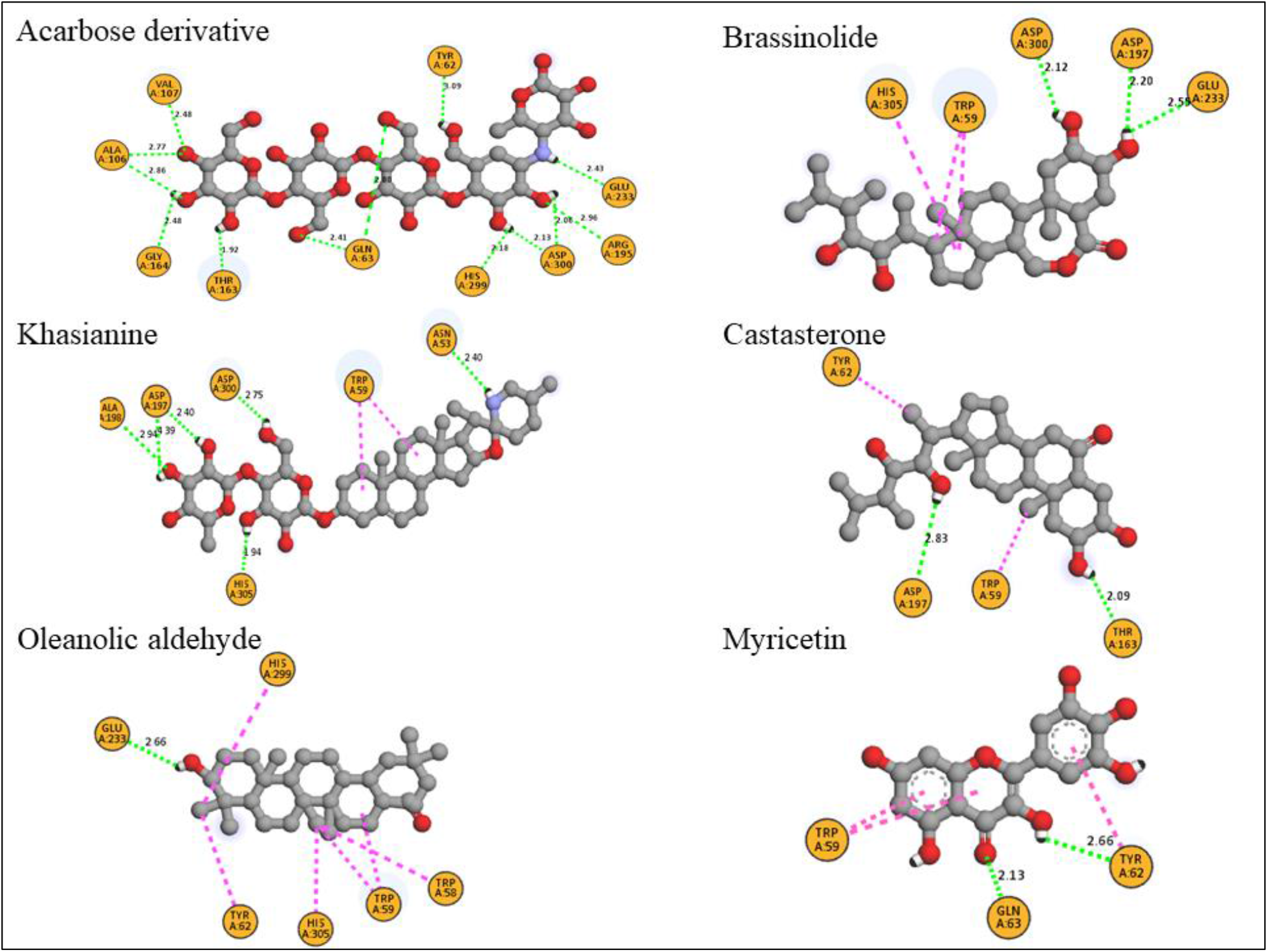
The interaction of human pancreatic α-amylase (HPA) with selected phytochemical compounds of *X. stuhlmannii* (Taub.) and acarbose. Hydrogen bonds are represented in green dotted lines. The purple dotted lines represent hydrophobic interactions (π-alkyl, π-σ, π-π stacking and π-π T-shaped).

**Figure 3.**
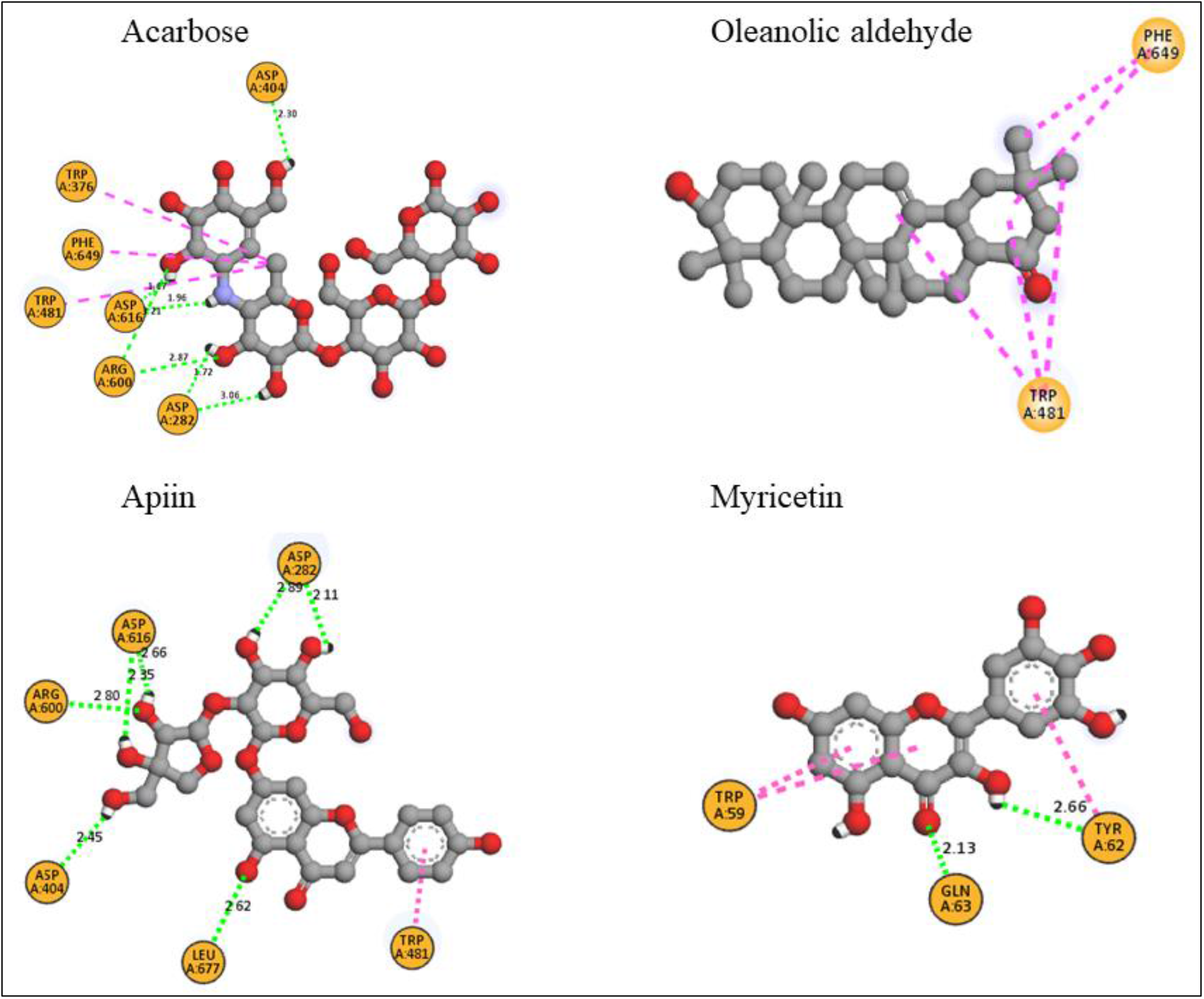
The interaction of human lysosomal acid-α-glucosidase (hGAA) with selected phytochemical compounds of *X. stuhlmannii* (Taub.) and acarbose. Hydrogen bonds are represented in green dotted lines. The purple dotted lines represent hydrophobic interactions (π-alkyl, π-σ, π-π stacking and π-π T-shaped).

**Table 3.** Hydrogen bonding (H-bonding) and binding affinity of selected phytochemical compounds against HPA and GAA enzymes.

| Enzyme | Phytochemical compound | Hydrogen bond (H-bond) interacting residues and the H-bond distance (Å) | Number of H-bonds | Binding affinity (kcalmol<sup>-1</sup>) |
| --- | --- | --- | --- | --- |
| HPA | Acarbose derivative | TYR62 (3.09), GLN63 (2.41), ALA106 (2.77), VAL107 (2.48), THR163 (1.92), GLY164 (2.48), ARG195 (2.96), GLU233 (2.43), HIS299 (2.18), ASP300 (2.06) | 10 | -9.3 |
|  | Khasianine | ASN53 (2.40), ASP197 (2.40), ALA198 (2.94), ASP300 (2.75), His305 (1.94) | 5 | -10.3 |
|  | Brassinolide | ASP197 (2.20), GLU233 (2.55), ASP300 (2.12) | 3 | -10.0 |
|  | Oleanolic aldehyde | GLU233 (2.66) | 1 | -9.4 |
|  | Castasterone | ASP197 (2.83), THR163(2.09) | 2 | -9.3 |
|  | Apiin | ASP197 (2.22), GLU233 (2.46), ILE235 (2.36) | 3 | -9.2 |
|  | Myricetin | TYR62 (2.66), GLN63 (2.13) | 2 | -9.0 |
|  | Ursolic acid | GLU233 (2.29), ASP300 (2.60) | 2 | -8.9 |
|  | Myricitin | GLN63 (2.39), ASP197 (2.97), | 2 | -8.5 |
| hGAA | Acarbose | ASP282 (1.72), ASP404 (2.30), ASP616 (1.67), ASP600 (2.87) | 4 | -8.1 |
|  | Oleanolic aldehyde | None |  | -8.4 |
|  | Apiin | ASP282 (2.11), ASP404 (2.45), ARG600 (2.80), ASP616 (2.35), LEU677 (2.62) | 5 | -8.3 |
|  | Khasianine | ARG281 (2.10), ASP616 (2.04) | 2 | -8.1 |
|  | Ursolic acid | ARG600 (2.28), ASP282 (2.13) | 2 | -7.8 |
|  | Castasterone | ASP282 (1.96), LEU677 (2.86), LEU678 (2.26) | 3 | -7.6 |
|  | Myricitin | ARG281 (2.70), ASP282 (2.52) | 2 | -7.4 |
|  | Myricetin | ASP404 (2.27), HIS674 (2.56) | 2 | -7.1 |

### The physicochemical, drug-likeness and ADME properties of Xeroderris stuhlmannii Taub. phytochemical compounds

The physicochemical, drug-likeness, bioavailability and medicinal properties of the selected compounds were evaluated using Swiss ADME, and are shown in Tables 4–7. Tables 4 and 3 show the physicochemical properties of the phytochemical compounds of *X. stuhlmannii* (Taub.). Only acarbose, khasianine and apiin had molecular weight greater than 500 g/mol. The number of heavy atoms ranged from 23-51 for all docked compounds, and only apiin, myricetin and myricitin had sixteen aromatic heavy atoms. All docked compounds had 10 or fewer rotatable bonds. In addition, five compounds including oleanolic aldehyde, ursolic acid and castasterone had 12 or fewer H-bond donors and acceptors (Table 4). The fraction of sp^3^ carbon atoms (Fsp^3^) was greater than 0.9 for all phytochemical chemicals studied except apiin (0.42), myricitin (0.29) and myricetin (0) (Table 4). The polarity index of the selected compounds was assessed by the topological surface area (TPSA) descriptor, and ranged from 37.3 to 225.1 (Table 4). The consensus LogP (a measure of solubility) of the docked compounds ranged from −6.22 to 6.32 (Table 5). Only acarbose (cLogP=−6.22) is highly soluble in water, whereas apiin (cLogP= −0.72), myricitin (cLogP= −0.23) and myricetin (cLogP= 0.79) are less soluble. Ursolic acid and oleanolic acid are highly insoluble (cLogP>5). The drug-likeness properties of the compounds showed that only brassinolide and castasterone did not violate Lipinski’s Rule of Five with a bioavailability score of 0.55 (Table 6). Oleanolic aldehyde, myricetin and ursolic acid violated one Lipinski’s Rule of Five with bioavailability scores of 0.55, 0.55 and 0.85, respectively. Myricitin, acarbose, apiin and khasianine had at least two Lipinski’s rule violations, and were predicted to have poor bioavailability. Only myricetin had good bioavailability and solubility properties with fewer violations of the Lipiski, Ghose, Veber, Egan and Muegge rules, and was best described as a druglike compound. Myricitin, and myricetin were flagged to have at least one PAINS substructure whereas, brassinolide, castasterone and apiin were not flagged to have a BRENK substructure (see Table 7). The predicted synthetic accessibility values of the phytochemical compounds are shown in Table 7, and ranged from 3.29 to 9.1. The synthetic accessibility values presented here demonstrate that *X. stuhlmannii* (Taub.) phytochemical compounds have moderate to complex synthetic route. Collectively, the results presented here indicate that myricetin has drug-likeness properties, and can be a good inhibitor of HPA and hGAA.

**Table 4.** Physicochemical properties of the phytochemical compounds of *X. stuhlmannii* (Taub.) determined using Swiss ADME.

| Phytochemical compound | Molecular Formula | Molecular Weight/gmol<sup>-1</sup> | #Heavy atoms | #Aromatic heavy atoms | #Rotatable bonds | #H-bond acceptors | #H-bond donors | Fraction Fsp<sup>3</sup> | MR | TPSA |
| --- | --- | --- | --- | --- | --- | --- | --- | --- | --- | --- |
| Acarbose | C<sub>25</sub>H<sub>43</sub>NO<sub>18</sub> | 645.6 | 44 | 0 | 9 | 19 | 14 | 0.92 | 136.7 | 321.2 |
| Brassinolide | C<sub>28</sub>H<sub>48</sub>O<sub>6</sub> | 480.68 | 34 | 0 | 5 | 6 | 4 | 0.96 | 133.7 | 107.2 |
| Khasianine | C<sub>39</sub>H<sub>63</sub>NO<sub>11</sub> | 721.92 | 51 | 0 | 5 | 12 | 7 | 0.95 | 190.8 | 179.6 |
| Oleanolic aldehyde | C<sub>30</sub>H<sub>48</sub>O<sub>2</sub> | 440.7 | 32 | 0 | 1 | 2 | 1 | 0.9 | 135.1 | 37.3 |
| castasterone | C<sub>28</sub>H<sub>48</sub>O<sub>5</sub> | 464.68 | 33 | 0 | 5 | 5 | 4 | 0.96 | 132.6 | 98.0 |
| Apiin | C<sub>26</sub>H<sub>28</sub>O<sub>14</sub> | 564.49 | 40 | 16 | 7 | 14 | 8 | 0.42 | 132.6 | 229.0 |
| Myricitin | C<sub>21</sub>H<sub>20</sub>O<sub>12</sub> | 464.38 | 33 | 16 | 3 | 12 | 8 | 0.29 | 111.0 | 210.5 |
| Myricetin | C<sub>15</sub>H<sub>10</sub>O<sub>8</sub> | 318.24 | 23 | 16 | 1 | 8 | 6 | 0 | 80.1 | 151.6 |
| Ursolic acid | C<sub>30</sub>H<sub>48</sub>O<sub>3</sub> | 456.7 | 33 | 0 | 1 | 3 | 2 | 0.9 | 136.9 | 57.5 |

**Table 5.**
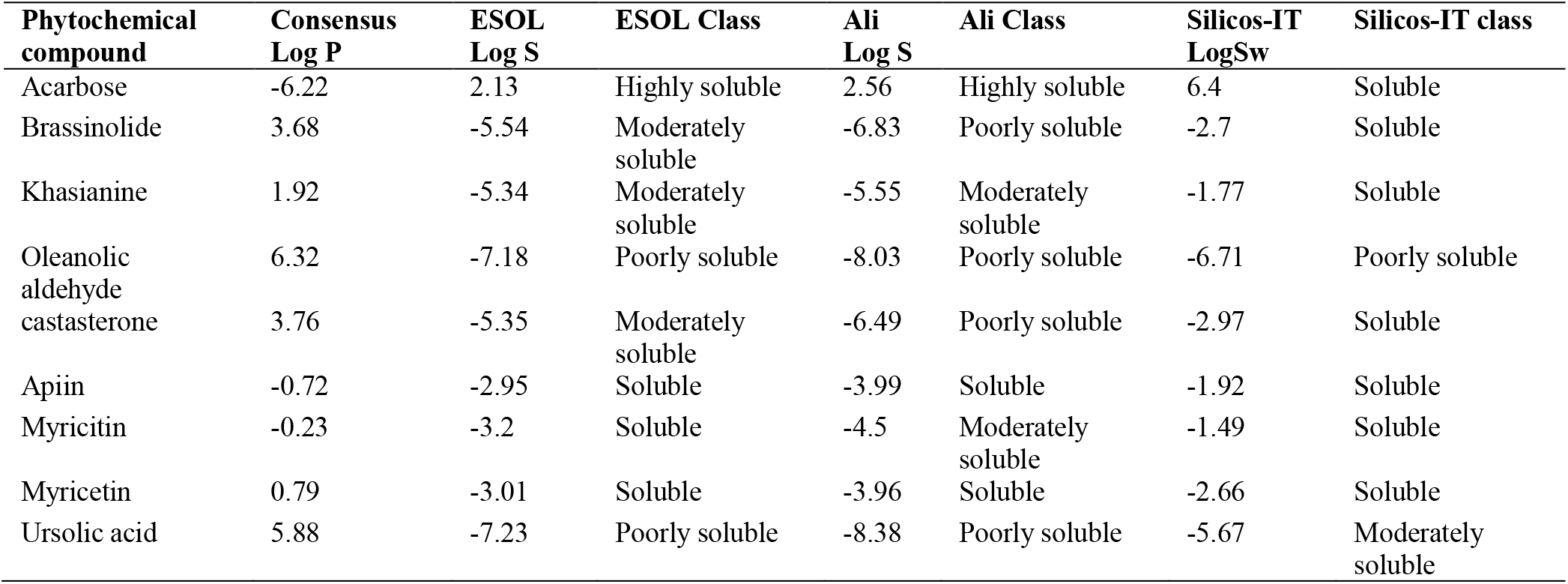
Lipophilicity and water solubility properties of the of the phytochemical compounds of *X. stuhlmannii* (Taub.) determined using Swiss ADME.

| Phytochemical compound | Consensus Log P | ESOL Log S | ESOL Class | Ali Log S | Ali Class | Silicos-IT LogSw | Silicos-IT class |
| --- | --- | --- | --- | --- | --- | --- | --- |
| Acarbose | -6.22 | 2.13 | Highly soluble | 2.56 | Highly soluble | 6.4 | Soluble |
| Brassinolide | 3.68 | -5.54 | Moderately soluble | -6.83 | Poorly soluble | -2.7 | Soluble |
| Khasianine | 1.92 | -5.34 | Moderately soluble | -5.55 | Moderately soluble | -1.77 | Soluble |
| Oleanolic aldehyde | 6.32 | -7.18 | Poorly soluble | -8.03 | Poorly soluble | -6.71 | Poorly soluble |
| castasterone | 3.76 | -5.35 | Moderately soluble | -6.49 | Poorly soluble | -2.97 | Soluble |
| Apiin | -0.72 | -2.95 | Soluble | -3.99 | Soluble | -1.92 | Soluble |
| Myricitin | -0.23 | -3.2 | Soluble | -4.5 | Moderately soluble | -1.49 | Soluble |
| Myricetin | 0.79 | -3.01 | Soluble | -3.96 | Soluble | -2.66 | Soluble |
| Ursolic acid | 5.88 | -7.23 | Poorly soluble | -8.38 | Poorly soluble | -5.67 | Moderately soluble |

**Table 6.** Drug-likeness properties and oral bioavailability of the phytochemical compounds of *X. stuhlmannii* (Taub.) determined using Swiss ADME.

| Phytochemical compound | Lipinski violations | Ghose violations | Veber violations | Egan Violations | Muegge violations | Bioavailability Score |
| --- | --- | --- | --- | --- | --- | --- |
| Acarbose | 3 | 4 | 1 | 1 | 5 | 0.17 |
| Brassinolide | 0 | 3 | 0 | 0 | 0 | 0.55 |
| Khasianine | 3 | 3 | 1 | 1 | 5 | 0.17 |
| Oleanolic aldehyde | 1 | 3 | 0 | 1 | 1 | 0.55 |
| castasterone | 0 | 2 | 0 | 0 | 0 | 0.55 |
| Apiin | 3 | 3 | 1 | 1 | 3 | 0.17 |
| Myricitin | 2 | 0 | 1 | 1 | 3 | 0.17 |
| Myricetin | 1 | 0 | 1 | 1 | 2 | 0.55 |
| Ursolic acid | 1 | 3 | 0 | 1 | 1 | 0.85 |

**Table 7.** Medicinal chemistry properties of the phytochemical compounds of *X. stuhlmannii* (Taub.) determined using Swiss ADME.

| Phytochemical compound | PAINS #alerts | Brenk #alerts | Leadlikeness #violations | Synthetic Accessibility |
| --- | --- | --- | --- | --- |
| Acarbose | 0 | 1 | 2 | 7.34 |
| Brassinolide | 0 | 0 | 2 | 6.14 |
| Khasianine | 0 | 1 | 1 | 9.1 |
| Oleanolic aldehyde | 0 | 2 | 2 | 5.99 |
| castasterone | 0 | 0 | 2 | 5.74 |
| Apiin | 0 | 0 | 1 | 6.08 |
| Myricitin | 1 | 1 | 1 | 5.32 |
| Myricetin | 1 | 1 | 0 | 3.27 |
| Ursolic acid | 0 | 1 | 2 | 6.21 |

To explore the pharmacokinetic properties (ADME) of the selected compounds, we computed the human intestinal absorption (HIA), caco-2 permeability (caco-2), P-glycoprotein (P-gp) inhibitor and substrate parameters in order to predict the absorption of the compounds in the gastrointestinal tract (Table 8). As shown in Table 5, khasianine, apiin, myricitin, and acarbose have low human intestinal absorption (HIA) probabilities. The computed caco-2 permeability parameters ranged from −4.85 cm/s to −6.35 cm/s. Caco-permeabilities greater than −5.15 are considered optimal (Xiong et al., 2021). Our results showed that only castasterone and brassinolide have better intestinal permeabilities compared to other phytochemical compounds. None of the compounds are considered as P-gp inhibitors but apiin and acarbose are categorized as P-gp substrates. Furthermore, we predicted that oleanolic aldehyde and ursolic acid are capable of crossing the blood brain barrier. Brassinolide, castasterone, castasterone and myricetin have low therapeutic index (>90% Plasma Protein Binding). The widespread microsomal cytochrome P450 (CYP) enzymes play an important role in phase 1 biotransformation of endogenous and exogenous compounds including many pharmaceutical drugs and phytochemical compounds (Guengerich, 2015). We therefore evaluated the inhibition of the major drug metabolizing CYPs by the phytochemical compounds, and showed that myricetin is an inhibitor of CYP1A2, while oleanolic aldehyde was predicted to be an inhibitor of CYP2D6 and CYP3A4. All other compounds were not classified as inhibitors of the CYP enzymes. Finally, castasterone and brassinolide have higher drug clearance than that of acarbose and other compounds. Overall, the ADME results showed none of the phytochemical compounds have a complete set of pharmacokinetic properties needed for their disposition in the blood stream.

**Table 8.** Pharmacokinetics properties of the phytochemical compounds of the *X. stuhlmannii* (Taub.) determined using ADMElab 2.0.

| Phytochemical compound | Human intestinal absorption (HIA) | Caco-permeability | Pgp inhibitor | Pgp substrate | PPB (%) | BBB penetration | CYP 1A2 inhibitor | CYP 2C19 inhibitor | CYP 2C9 inhibitor | CYP 2D6 inhibitor | CYP 3A4 inhibitor | CL (mL/min/kg) |
| --- | --- | --- | --- | --- | --- | --- | --- | --- | --- | --- | --- | --- |
| Acarbose | 1.0 | -6.35 | 0 | 0.85 | 8.2 | 0.39 | No | No | No | No | No | 0.37 |
| Brassinolide | 0.13 | -4.85 | 0.1 | 0.06 | 92.7 | 0.23 | No | No | No | No | No | 19.35 |
| Khasianine | 0.92 | -5.36 | 0 | 0.01 | 81.4 | 0.03 | No | No | No | No | No | 0.94 |
| Oleanolic aldehyde | 0.01 | -5.09 | 0.03 | 0 | 80.7 | 0.93 | No | No | No | Yes | Yes | 6.24 |
| Castasterone | 0.27 | -4.85 | 0.02 | 0.05 | 93.9 | 0.56 | No | No | No | No | No | 21.38 |
| Apiin | 0.96 | -6.27 | 0 | 0.95 | 81.4 | 0.16 | No | No | No | No | No | 1.66 |
| Myricitin | 0.70 | -6.27 | 0 | 0.58 | 87.7 | 0.01 | No | No | No | No | No | 5.26 |
| Myricetin | 0.04 | -5.65 | 0 | 0.01 | 92.8 | 0.01 | Yes | No | No | No | No | 7.72 |
| Ursolic acid | 0.01 | -5.22 | 0 | 0 | 98.8 | 0.72 | No | No | No | No | No | 3.67 |

## Discussion

It is important to recognize that bioactive phytochemical compounds such as flavonoids, terpenoids, saponins, polyphenols, alkaloids and glycosides play an important role in human health. For example, polyphenolic compounds such as myricetin possess antioxidant activity, and can scavenge highly reactive free radicals within the biological system (Barzegar, 2016). In this study, we determined the antioxidant activity of *X. stuhlmannii* (Taub.) root and bark extracts against DPPH radicals, and showed that the root and bark extracts significantly scavenged DPPH radicals with low IC_50_ values (<0.013 μg/mL). The antioxidant activities of the extracts are likely attributed to the bioactive polyphenolic and flavonoid compounds in the crude extracts. In addition, we showed that ethyl acetate extracted more polyphenols and flavonoids than methanol. This observation is in agreement with published results that showed that ethyl acetate fractions are rich in flavonoids (Bhatia et al., 2019).

Inhibition of the activity of enzymes involved in carbohydrate metabolism including α-glucosidase and α-amylase is one of the novel approaches developed to treat T2DM mellitus and its complications. Novel α-glucosidase inhibitors delay the overall digestion of carbohydrates by increasing the digestion period and reducing the rate of intestinal glucose absorption, which in turn diminishes postprandial hyperglycemia (Mohammed and Tajuddeen, 2022). Here we investigated the *in vitro* inhibitory properties of *X. stuhlmannii* (Taub.) extracts on α-amylase and α-glucosidase enzymes because it is routinely used to treat many ailments including T2DM without proper scientific evidence to support its use (Karou et al., 2011). Our results showed that the half maximal inhibitory concentration (IC_50_) of the extracts were significantly lower than that of acarbose. The high potency of the extracts (low IC_50_ values) compared to acarbose against digestive enzymes is likely due to the presence of bioactive polyphenols and flavonoid compounds acting alone or in combination to inhibit the digestive enzymes (Selemani et al., 2021).

Molecular docking remains one of the most popular *in silico* approaches used in drug discovery to virtually screen for hit compounds in virtual libraries containing millions of molecular structures against a variety of drug targets with known three-dimensional structures (Gentile et al., 2020). The molecular modelling approach provides important information on the ligands’ binding affinity and can effectively predict different binding modes of the ligand in the active site of the target molecule (Meng et al., 2011). Molecular docking approaches have successfully been utilized to identify novel natural product inhibitors of human α-amylase (HPA) and human α-glucosidase (hGAA) (Mugaranja and Kulal, 2022). Here, we used Autodock Vina (Scripps Research, San Diego) to identify novel photochemical compounds of *X. stuhlmannii* (Taub.) extracts that may inhibit digestive enzymes. Among the thirty-six docked phytochemical compounds, only five (Khasianine, brassinolide, oleanolic aldehyde and apiin) had higher docking scores compared to acarbose, and were therefore predicted to inhibit α-glucosidases. These compounds inhibited digestive enzymes by forming strong intermolecular forces including H-bonds with active site residues. The inhibitory effects of the phytochemical ligands are consistent with the low IC_50_ values of the extracts. In addition, we include myricetin, myricitrin and ursolic acid in molecular docking studies because they are known to inhibit α-glucosidase enzymes (Castro et al., 2015; Kim et al., 2020; Williams et al., 2012).

Within the active site of HPA, three essential acidic residues (ASP197, GLU233 and ASP300) exist that catalyze the breakdown of glyosidic bonds (Maurus et al., 2008; Williams et al., 2012). Inhibitors that form strong H-bonds with these acidic residues can delay carbohydrate hydrolysis, and can ultimately reduce postprandial hyperglycemia. Our studies showed that all docked phytochemical compounds, with the exception of myricetin can form at least one H-bond with the catalytic residues. A closer look at the active site of hGAA showed that the catalytic nucleophile and acid/base residues are ASP518 and ASP616 (Roig-Zamboni et al., 2017). We redocked acarbose in the active site of hGAA, together with eight other selected phytochemical compounds of *X. stuhlmannii* (Taub.). Only acarbose, apiin and khasianine formed a H-bond with the catalytic residue, ASP616. In addition, our results showed that apiin formed more H-bonds than other phytochemical compounds, and can be a novel inhibitor of hGAA. However, to the best of our knowledge the inhibitory effect of apiin on digestive enzymes have not yet been explored.

Evaluation of pharmacokinetic and pharmacodynamic properties of hits in drug discovery is important because it helps us understand how drugs behave in the body and how the body reacts to certain drugs. Pharmacokinetic prediction studies presented in this work showed that none of the hit phytochemical compounds exhibited all ADME properties. Consistent with published results, our pharmacokinetic prediction studies also showed that myricetin had fewer violations than other compounds, and can be a novel inhibitor of α-glucosidases. The antidiabetic activity of myricetin and its derivatives has been reported and a high-resolution X-ray crystal structure of human pancreatic α-amylase (HPA) complexed with myricetin has been solved (Williams et al., 2012). Myricetin binds to the active site and interacts directly with catalytic residues of HPA and reduces the normal conformational flexibility of the substrate binding cleft (Fu et al., 2021; Williams et al., 2012). The antidiabetic role of ursolic acid is mediated through insulin secretion and insulinomimetic effect on glucose uptake, synthesis and translocation of GLUT4 by a mechanism of cross-talk between calcium and protein kinases (Castro et al., 2015). However, our results showed that ursolic acid is highly insoluble in aqueous and lipid environments, and may require structural modifications to improve its solubility. Myricitrin improves T2DM by significantly decreasing the fasting blood glucose levels, improving glucose intolerance and increasing pancreatic β-cell mass (Kim et al., 2020).

## Conclusions

In this work, we identified using docking studies novel α-amylase and α-glucosidase inhibitors of *X. stuhlmannii* (Taub.) capable of delaying carbohydrate metabolism by inhibiting α-glucosidase enzymes. These compounds may inhibit carbohydrate metabolism as individuals or in combination by competitively binding to the active site of the enzymes, and preventing the substrates from accessing the active site. The inhibitory effect of the compounds on the digestive enzymes likely contributed to their low IC_50_ values. Molecular docking and pharmacokinetic prediction studies showed myricetin may be a novel inhibitor of α-glucosidases capable of alleviating T2DM symptoms by inhibiting these enzymes.

## Author Contributions

FR, EM and GM were involved in concept development. BN, CVPM and EM performed the *in vitro* experiments, and JTB performed the in-silico modeling experiments. FR, GM, JTB, and BN wrote the manuscript. All authors have read and agreed to the published version of the manuscript.

